# CST interacts with the cohesin complex and promotes chromosome cohesion

**DOI:** 10.1101/2021.04.11.439363

**Authors:** P. Logan Schuck, Jason A. Stewart

## Abstract

Sister chromatid cohesion (SCC) is established during DNA replication by loading of the cohesin complex on newly replicated chromatids. Cohesin must then be maintained until mitosis to prevent segregation defects and aneuploidy. How SCC is established and maintained until mitosis remains incompletely understood and emerging evidence suggests that replication stress can lead to premature SCC loss. Here, we report that the single-stranded DNA-binding protein CTC1-STN1-TEN1 (CST) aids in SCC. CST primarily functions in telomere length regulation but also has known roles in replication restart and DNA repair. Following depletion of CST subunits, we observed an increase in the complete loss of SCC. Additionally, we determined that CST interacts with the cohesin complex. Unexpectedly, we did not find evidence of defective cohesion establishment or mitotic progression in the absence of CST. However, we did find that treatment with various replication inhibitors increased the association between CST and cohesin. Since replication stress was recently shown to induce SCC loss, we supposed that CST may be required to maintain SCC following fork stalling. In agreement with this idea, SCC loss was greatly increased in CST-depleted cells following exogenous replication stress. Based on our findings, we propose that CST aids in the maintenance of SCC at stalled replication forks to prevent premature cohesion loss.

## Introduction

As DNA is replicated, the sister chromatids must be held together until mitosis to ensure chromosomes are properly segregated between daughter cells. This process, known as sister chromatid cohesion (SCC), is facilitated by the cohesin complex (1,2). In mammals, the cohesin complex is composed of a ring-like structure that encircles the DNA. The core structure is composed of SMC3, SMC1A, SCC1 (also known was RAD21) and SA1 or SA2. Cohesin loading and removal is tightly regulated during the cell cycle (3-5). In G1 and early S-phase, cohesin is loaded onto chromatin but is not stably bound. As the DNA is replicated, cohesin is passaged to the replicated sister chromatids and becomes stably bound until mitosis when it is removed to allow segregation of the chromatids into daughter cells. In addition to its essential role in SCC, the cohesin complex is involved in organizing topologically associated domains for cellular processes such as DNA repair and gene expression (6). Cohesin has also been implicated in replication fork restart with perturbations in cohesion dynamics following fork stalling promoting premature SCC loss (7-9).

In this study, we present unexpected findings that human CTC1-STN1-TEN1 (CST) helps to maintain SCC. CST is an RPA-like, ssDNA binding protein that is conserved from yeast to humans (10). Like RPA, CST contains multiple oligosaccharide-oligonucleotide folds (OB-folds) that allow dynamic binding to the DNA (11,12). CST primarily functions in telomere length regulation, however, it has also been shown to function in DNA replication and repair (13,14). While its role in DNA replication is still not well understood, it is proposed to aid in the replication of G-rich DNA sequences, such as telomeres, promote dormant origin firing and negatively regulate origin licensing (15-18). CST also interacts with several components of the replication machinery, including DNA polymerase α-primase, the MCM2-7 helicase and AND-1/Ctf4 (16,19,20). Together, these findings provide strong evidence that CST is involved in DNA replication, presumably as a specialized versus general replication factor.

Here, we report that depletion of CST leads to premature SCC loss. Furthermore, we show that CST interacts with the cohesin subunits, SMC1A and SMC3, suggesting that CST may directly influence SCC. Upon further investigation, we found that the interaction between cohesin and CST is increased following replication stress and that CST prevents SCC loss following treatment with several replication inhibitors. Together, our findings identify CST as a new factor involved in preventing premature cohesion loss and suggest that it does so by stabilizing chromosome cohesion following replication fork stalling.

## Results

### Depletion of CST results in sister chromatid cohesion loss

While performing telomere-FISH in STN1 knockdown cells, we consistently observed metaphase spreads that had lost SCC. To determine whether depletion of STN1 increased SCC loss, we quantified the number of metaphase spreads with SCC loss in HeLa cells with stable shRNA knockdown of STN1 (shSTN1) (Fig 1A-B) (18). In agreement with our observation, we observed a 2-to 4-fold increase in premature SCC loss in two separate shSTN1 clones, shSTN1-6 and shSTN1-7 (Fig 1C). Furthermore, this increase was largely rescued by stable expression of a Flag-tagged shRNA-resistant STN1 construct in shSTN1-7 cells (shSTN1-7 +Flag-STN1). Previous studies suggest that a common off-target effect of RNA interference is MAD2 depletion (21). Loss of MAD2 also results in premature SCC loss. MAD2 levels were measured in the shSTN1 cells and no significant changes were observed compared to controls (Fig. 1A).

**Figure 1.**
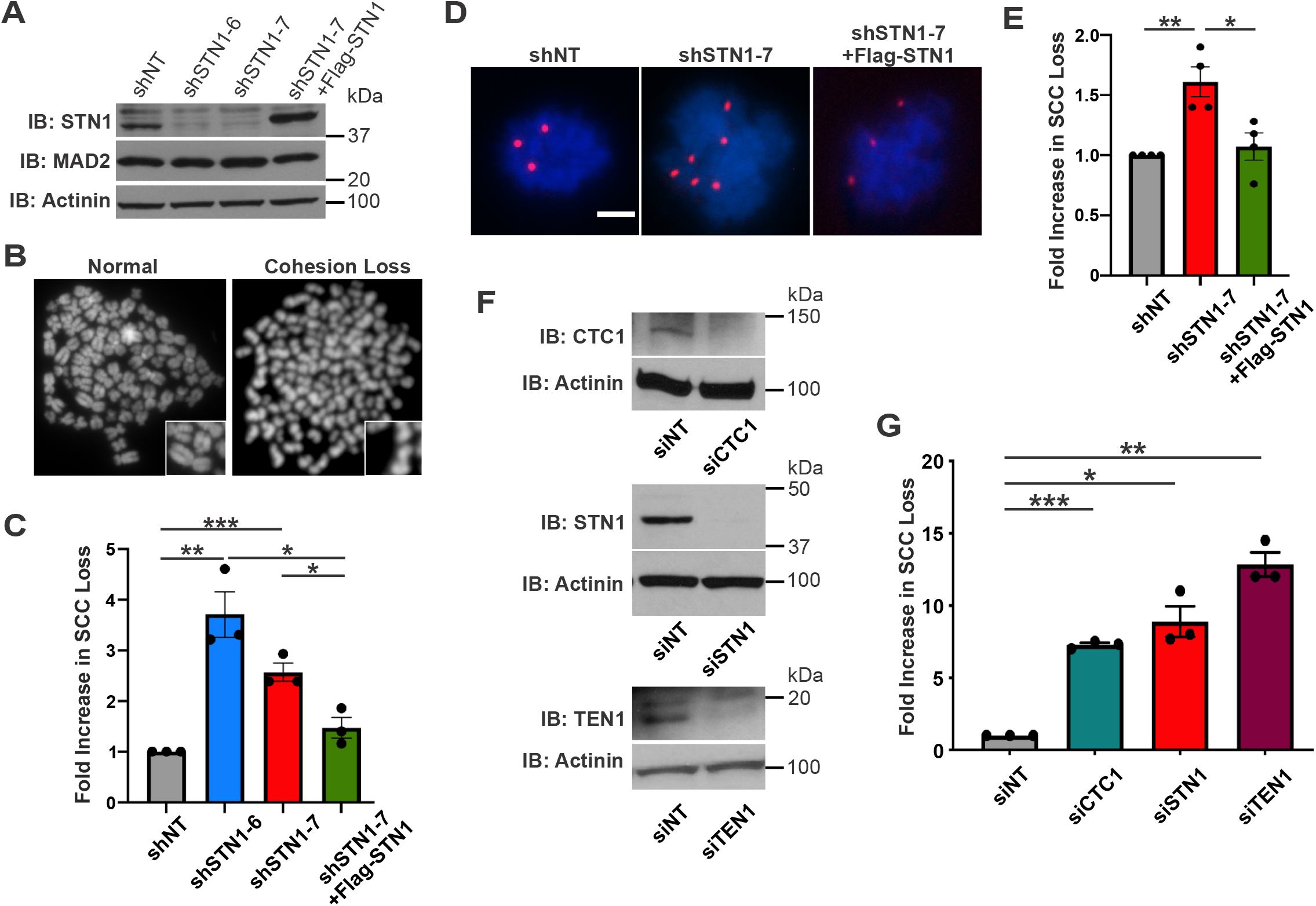
CST deficiency results in SCC loss. (A) Western blot analysis of STN1 knockdown and MAD2 levels in HeLa cells. Actinin: loading control. shNT: non-targeting shRNA; shSTN1: shRNA to STN1; and shSTN1 +Flag-STN1: shSTN1-7 cells plus shRNA-resistant Flag-STN1. (B) Example images of metaphase spreads with normal chromosomes or SCC loss. (C) Fold increase in cohesion loss following metaphase spread analysis, as indicated. n=3 independent, biological replicates. (D) Representative images of chromosome FISH from cells isolated by mitotic shake-off. Red: centromere 6 probe, Blue: DAPI. Scale bar: 5 μm (E) Fold increase in nuclei with >4 chromosome 6 foci, indicating SCC loss. n=4 independent, biological replicates. (F) Knockdown of CTC1, STN1 or TEN1 by siRNA in HeLa cells. siNT: non-target control, Actinin: loading control. (G) Graph of SCC loss following metaphase spread analysis, as indicated. n=3 independent, biological replicates. *P*-values were calculated by an unpaired, two-sided t-test (*P < 0.05, **P < 0.01, ***P < 0.001).

To confirm our findings, we next performed mitotic shake-off to measure SCC loss following STN1 depletion. Processing of cells for standard metaphase spread analysis includes treatment with a hypotonic solution, which can release proteins from the chromatin (22). If cohesion is only partially lost, this can exacerbate the number of cells with SCC loss. To address this possibility, mitotic cells were collected, fixed and spun onto slides without hypotonic treatment. Unlike standard metaphase spread preparation, cytospun metaphase cells are not clearly separated. Therefore, a chromosome specific probe to centromere six was used to assess SCC loss by fluorescence *in situ* hybribization, as previously described (23). Since HeLa cells are triploid for chromosome six, three spots indicate the retention of SCC, whereas greater than three spots indicate SCC loss. Consistent with metaphase spread analysis, knockdown of STN1 lead to a significant increase in SCC loss (Fig. 1D-E). However, the fold increase was less than that observed by standard metaphase spread analysis (Fig. 1C). This may be due to weakened but not complete SCC loss in a subset of cells or differences between the methodologies used.

Next, we determined whether cohesion loss was specific to STN1 depletion or due to a general loss of CST. siRNA knockdown of CTC1, STN1 or TEN1 was performed followed by metaphase spread analysis (Fig 1F-G). Like shSTN1 cells, transient siRNA knockdown of individual CST subunits resulted in increased SCC loss. Finally, we confirmed that this phenotype was not cell type specific by measuring SCC loss in HCT116 cells with conditional CTC1 knockout, which also showed increased SCC loss (Fig. S1). Together, these results indicate that CST promotes SCC.

### CST interacts with the cohesin complex

We next addressed whether CST interacts with the cohesin complex by co-immunoprecipitation (co-IP) and proximity ligation assay (PLA) (Fig 2). IP of epitope-tagged CST pulled down both endogenous SMC3 and SMC1A (Fig 2A). STN1 alone also showed interaction with SMC3, although at lower levels. Pulldown of CTC1 showed little to no association with SMC3 or SMC1A. This suggests that the entire CST complex, or at least multiple subunits, may be required for stable interaction. As further confirmation of CST-cohesin association, we identified both SMC1A and SMC3 by mass spectrometry after IP of Flag-STN1 in cells overexpressing all three CST subunits (Fig S2).

**Figure 2.**
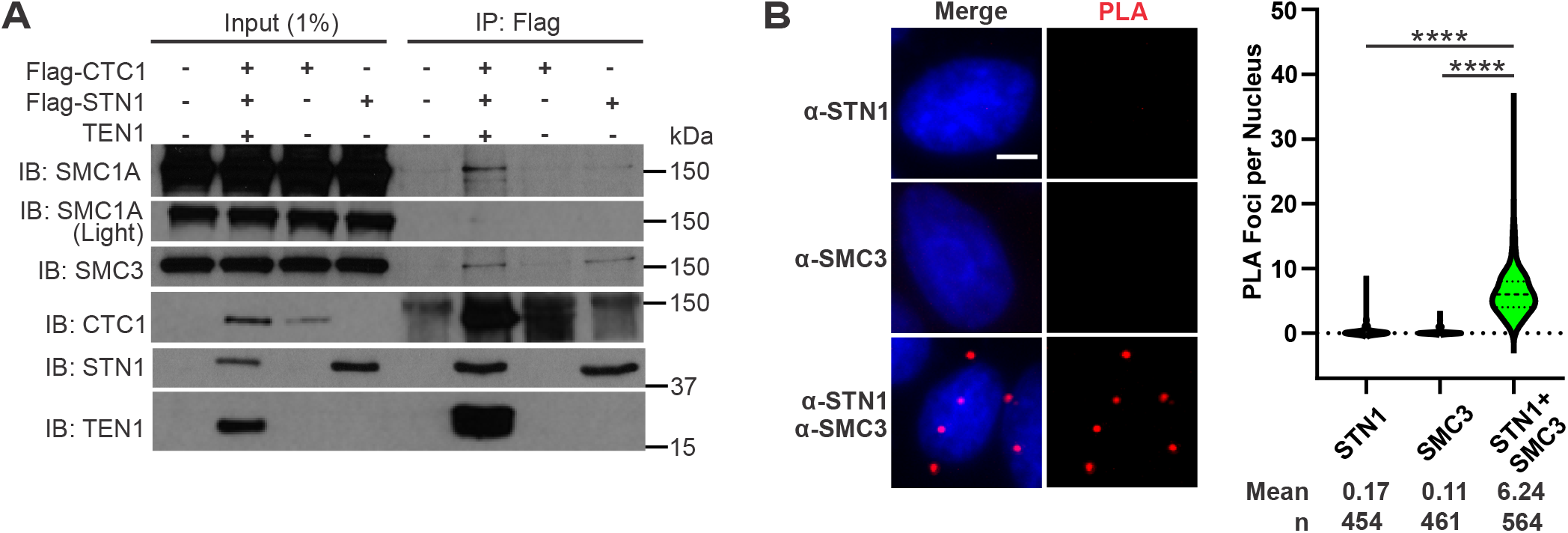
CST interacts with the cohesin complex. (A) Co-IP of nuclease-treated lysates from HEK 293T cells expressing Flag-tagged CTC1, Flag-tagged STN1 or the full CST complex. (B) Representative images of proximity ligation assay (PLA) performed in HeLa cells with antibodies to STN1 (α-STN1) or SMC3 (α-SMC3) alone or combined (α-STN1/α-SMC3). Red: PLA foci, Blue: DAPI. Scale bar: 5 μm (C) Violin plot of PLA foci per nucleus. Results are representative of four independent, biological experiments. Bold dashed line: median, dashed lines: first and third quartiles. *P*-values were calculated by an unpaired, two-sided t-test (****P < 0.0001).

To determine whether this interaction was also present in cells, PLA was performed using antibodies to endogenous STN1 and SMC3 (Figure 2B). PLA detects interactions through use of secondary antibodies with a segment of DNA attached. When these antibodies are in close proximity (<40 nm) the DNA segments are joined and the DNA is then amplified by rolling circle amplification and detected using fluorescent probes (24). PLA analysis with antibodies to endogenous STN1 and SMC3 showed ∼6 foci on average compared to single antibodies controls (Fig 2B). Combined with the co-IP data, these findings reveal an unanticipated interaction between CST and the cohesin complex.

### CST is not required for SMC3 acetylation, cohesin loading or mitotic progression

A possible explanation for SCC loss following CST depletion is that CST stabilizes cohesin loading or establishment (25,26). Prior to genome duplication, cohesin binding is unstable and cohesin molecules quickly associate/dissociate from the DNA (27). As the genome is duplicated, cohesin is transferred from the unreplicated to the replicated sister chromatids and becomes stably bound. This process is partially facilitated by the acetylation of SMC3 (Ac-SMC3) (28-31). Since CST aids in DNA replication, we tested whether Ac-SMC3 or the levels of total or chromatin-bound cohesin were decreased in STN1-depleted cells (Fig. 3A-C, S3A-B). However, we did not observe any changes in either Ac-SMC3 or chromatin-bound cohesin.

**Figure 3.**
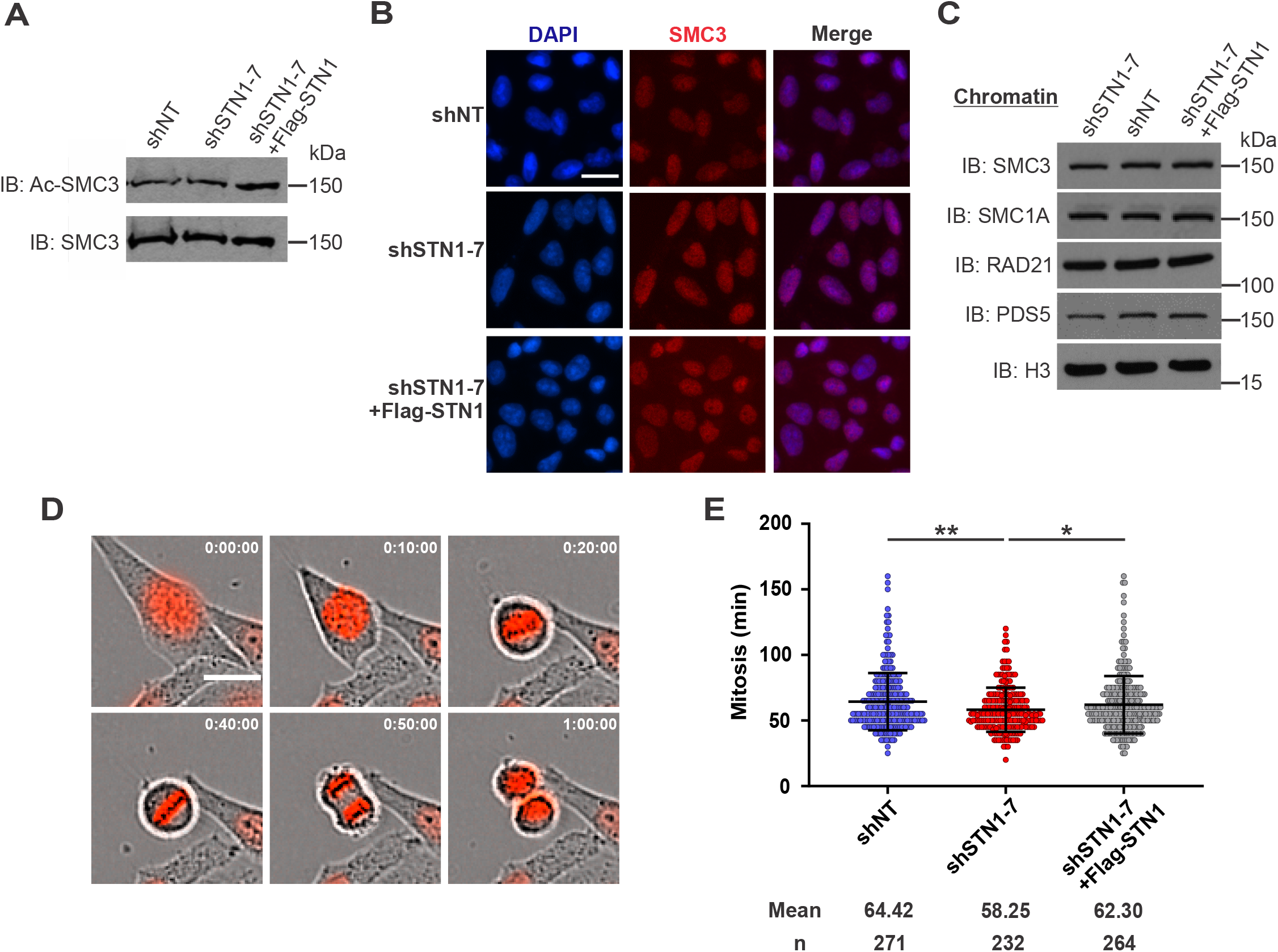
STN1 depletion does not affect cohesin levels or mitotic timing. (A) Western blot analysis of total SMC and acetylated-SMC3 in HeLa cells, as indicated. (B) Representative images of SMC3 levels in pre-extracted cells. DAPI: blue, SMC3: red. Scale bar: 20 μm (C) Western blot of chromatin-bound cohesin subunits. Histone H3 serves as a loading control. (D) HeLa cells stably expressing H2B-RFP were imaged in 5 min intervals for 3 hours. Scale bar: 20 μm (E) Dot plot of the time between nuclear envelope breakdown until cytokinesis in individual cells. Black lines and numbers below the graph indicate the average minutes. Error bars indicate the ±SEM of three independent biological experiments. n indicates the number of total cells scored. *P*-values were calculated by an unpaired, two-sided t-test (*P < 0.05, **P < 0.01).

Since our data suggest that cohesion establishment is unaffected by STN1 depletion, we investigated whether mitotic progression was altered using live cell imaging (Fig 3D-E). To visualize mitotic events, an H2B construct fused with red fluorescence protein was stably transduced into STN1-depleted or control cells (32). Cells were plated and imaged at 5 min intervals over a 3-hour period. The time from prophase to the completion of cytokinesis was then measured as a readout of mitotic progression (Figure 3D). Mitosis took approximately one hour in the HeLa cells, consistent with previous studies (33). On average, shSTN1 cells took ∼5 minutes less to complete mitosis compared to control cells (Fig. 3E). Further breakdown of the timing from prophase to metaphase or metaphase to cytokinesis did not reveal any significant changes (Fig S3C). While there was a slight decrease in overall mitotic progression in shSTN1 cells, these changes are unlikely to explain the increase in SCC loss. Why premature loss of SCC does not lead to defects in mitotic progression in cells lacking CST is not entirely clear. However, work in *Drosophila* suggest that premature SCC loss is not always sufficient to trigger a robust spindle assembly checkpoint and that mitosis can still occur (34,35). Moreover, recent work showed that various cancer cell lines continue to grow despite significant levels of cohesion loss (8,26). Accordingly, mitotic progression and cell division may be unaffected despite increased SCC loss in STN1 depleted cells.

### CST promotes SCC following replication stress

Since CST aids in DNA replication restart and SCC loss has been linked to replication stress, we sought to determine whether chemically induced replication stress would increase the interaction between CST and cohesin. Cells were treated with hydroxyurea (HU), aphidicolin (APH) or camptothecin (CPT). PLA was then performed with antibodies to endogenous STN1 and SMC3 (Fig. 4A). In all cases, we observed an ∼2-fold increase in PLA foci, indicating that cohesin-CST interaction is increased following replication stress compared to untreated cells. This suggested to us that CST may prevent SCC loss following replication stress, so we tested whether SCC loss was increased after treatment with replication inhibitors in shSTN1 cells. Following treatment with replication inhibitors, cells were released into fresh media for 6 hours to allow cells in S-phase to reach mitosis. They were then treated with colcemid, collected and prepared for metaphase spread analysis (Fig 4B). In agreement with previous findings, SCC loss was increased in the control cells when treated with the replication inhibitors (8,9). (CPT likely has the greatest effect because it is not reversible like HU and APH.) However, in the STN1 depleted cells, premature SCC loss was greatly increased above the shNT cells, consistent with CST promoting SCC following replication stress.

**Figure 4.**
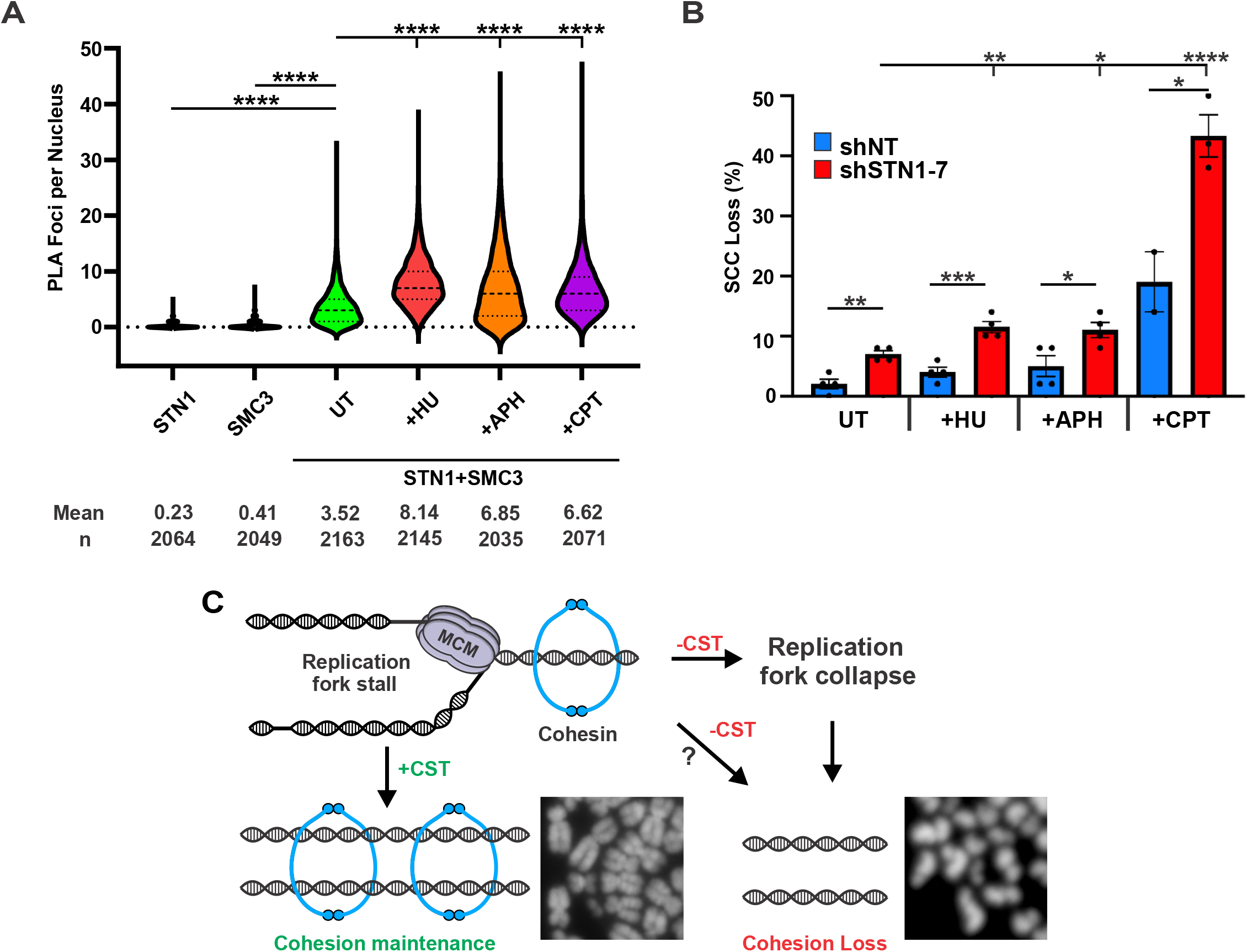
Replication inhibition increases the CST-cohesin interaction and SCC loss in STN1-depleted cells. (A) Violin plot of nuclear PLA foci in HeLa cells after treatment with DNA replication inhibitors. Drug treatments were performed for 2 hours for hydroxyurea (HU) and aphidicolin (APH). Camptothecin (CPT) treatment was performed for 1 hour. Bold dashed line: median, dashed lines: first and third quartiles. n=3 independent, biological replicates. (B) Cohesion loss in HeLa shNT and shSTN1 cells following replication stress. Cells were treated with replication inhibitors as in (A) and then released for 8 hours prior to metaphase spread preparation. Colcemid was added 2 hours prior to collection. n=3 independent, biological replicates. *P*-values were calculated by an unpaired, two-sided t-test (*P < 0.05, **P < 0.01, ***P < 0.001).

## Discussion

In this study, we present data that CST is involved in SCC and interacts with the cohesin complex. Despite significant levels of SCC loss in CST depleted cells, we were unable to detect any defects in cohesion establishment or mitotic progression. Instead, CST appears to be involved in SCC maintenance/re-establishment at stalled replication forks. This idea is in line with recent work demonstrating that replication stress, caused by expression of oncogenes or treatment with DNA replication inhibitors, induces SCC loss (8,9). Based on our data, we propose that CST acts at stalled replication forks to maintain or re-establish SCC following fork restart (Fig 4C). These findings highlight an unexpected function of CST in preserving genome integrity through the maintenance of SCC.

Over a decade ago, depletion of the replication fork stability factors Tipin/Tim and AND1 were shown to increase SCC loss, indirectly associating replication defects with chromatid cohesion (36-39). Additional studies have implicated other DNA replication and repair factors in chromatid cohesion and this past year a pair of studies provided direct evidence that chemically induced replication stress leads to SCC loss (8,9). These recent studies both suggest that the cohesin antagonist WAPL is involved in cohesion maintenance following replication stress. Benedict et al. propose a model where WAPL removes cohesin to allow replication fork restart through RAD51-dependent mechanisms. However, other studies, in both budding yeast and humans, suggest that cohesin may be necessary for fork restart or protection (7,40,41). Regardless of the mechanism, a picture is emerging where local cohesin architecture may be remodeled at stalled replication forks to allow restart. Interestingly, CST is proposed to load RAD51 following fork stalling (17). Therefore, CST could be a key player in cohesin dynamics at stalled forks by facilitating fork restart and cohesion reestablishment.

Another potential link between CST and cohesion maintenance is that CST interacts with both MCM2-7 and AND-1 (16). Recent work identified interaction between the MCM2-7 helicase and ESCO2 as necessary for cohesion establishment (25,28,31). As mentioned above, AND-1 is involved in replication fork stability and helps maintain SCC (38). Studies in yeast suggest that AND-1 (known as Ctf4 in yeast) stabilizes critical interactions between replisome components and the Chl1 helicase, which is involved in chromatid cohesion (42,43). We previously showed that loss of CST leads to decreased chromatin-bound AND-1, which could also provide a potential explanation for increased SCC loss in the absence of CST. We propose that interactions between CST, MCM2-7 and AND-1 are necessary for CST to help stabilize cohesion at stalled forks. Future studies will investigate how these interactions affect chromosome cohesion and fork restart/protection following fork stalling.

While the function of CST in replication restart/rescue is still unclear, CST is not a general replication factor but rather plays a specialized role in facilitating replication through G-rich regions of the genome (e.g. telomeres, CpG islands) (15,17,18). Since CST promotes replication at specific sites, one might predict that the absence of CST would cause partial versus total loss of chromosome cohesion, as has been previously observed with depletion of the cohesin subunits SA1 or SA2 (23). Instead, CST depletion leads to complete SCC loss by metaphase spread analysis and mitotic shake-off (Figure 1). These findings demonstrate that SCC loss is not restricted to telomeres or specific regions of the genome but instead a complete breakdown of SCC. The reason for complete cohesion loss remains unclear. However, recent studies suggest that complete cohesion loss is common across a variety of cancer cells, which seems to have little effect on cellular division (8). It is possible that the gradual accumulation of SCC loss due to replication stress or excessive DNA damage, triggers genome-wide cohesin unloading through an unknown mechanism. Perhaps such a pathway is used to induce cell death to prevent the propagation of cells with high levels of genome instability induced by replication stress. In cancer, such pathways could be disengaged and allow cell division and aneuploidy, despite SCC loss. Future studies will be needed to fully investigate the connection between replication stress, SCC and aneuploidy.

### Experimental procedures

#### Cell culture

HeLa 1.2.11 cells were cultured in RPMI 1640 media, HEK293T in DMEM, and HCT116 in McCoy’s 5A media at 37°C with 5% CO_2_. All media was supplemented with 10% fetal bovine serum and 1% penicillin/streptomycin. Stable HeLa 1.2.11 shRNA knockdown and HCT116 CTC1 inducible knockout lines have been previously described (18,44). Cell lines were regularly checked for mycoplasma contamination. siRNA experiments were performed using, 25 nM ON-TARGETplus siRNA SMART pools (Dharmacon) to CTC1 (L-014585-01), STN1 (L-016208-02), TEN1 (L-187549-00), or Non-targeting control (D-00180-10). siRNAs were transfected into cells with Lipofectamine RNAiMAX (Thermo Fisher Scientific) for 72 hours prior to collection.

#### Metaphase spreads

Metaphase spreads were prepared as previously described (45) and then stained for 10 min with Giemsa stain (Ricca Chemical). The slides were blinded prior to counting and at least 50 metaphase spreads were scored per slide per condition.

#### Whole cell lysate, chromatin fractionation and Western blot analysis

These techniques were performed, as previously described (16).

#### Antibodies and chemical inhibitors

##### Primary

SMC1A (Bethyl, A300-055A), SMC3 (Bethyl, A300-060A), acetylated-SMC3 (kindly provided by Dr. Prasad Jallepalli), RAD21 (SCC1) (Bethyl, A300-080A), SCC-112 (PDS5) (Bethyl, A300-089A), OBFC1 (STN1) (Novus, NBP2-01006), CTC1 (46), α-tubulin (Sigma, T-9026), TEN1 (47), MAD2 (Bethyl, A300-301A), α-Actinin (Santa Cruz, SC17829), H3 (Cell Signaling, 9715), Flag (Thermo Fisher, MA1-91878, PA1-984B)

##### Secondary

Thermo Fisher Scientific: anti-rabbit-HRP (Thermo Fisher, 32460), anti-mouse-HRP (Thermo Fisher, 32430), goat anti-rabbit Alexa Fluor 594 (Thermo Fisher, A-11037)

##### Chemical inhibitors

Cells were treated with aphidicolin (1 µM, Millipore-Sigma 178273) or hydroxyurea (2 mM, Millipore-Sigma 400046) for 2 hour or (S)-(+)-camptothecin (1 µM, Millipore-Sigma C9911) for 1 hour.

#### Live cell imaging

MSCV-H2B-mRFP1 was created by replacing GFP in MSCV-GFP with the H2B-mRFP1 from pCS-H2B-mRFP1, using NotI and XhoI. pCS-H2B-mRFP1 was a gift from Dr. Sean Megason (Addgene #53745) and MSCV-GFP was a gift from Dr. Tannishtha Reya (Addgene #20672). HeLa cells were transduced with retrovirus produced in HEK 293T and RFP positive cells were selected through two rounds of fluorescence-activated cell sorting. Approximately 3,000 cells were plated into 96 well IncuCyte ImageLock plates 24 h prior to imaging. Cells were then imaged at 5 min intervals for 3 h under a 20x objective, using the IncuCyte S3 Live-Cell Analysis System.

#### Fluorescent in-situ hybridization (FISH)

Following collection, cells were washed in 5 ml 1x PBS and then fixed by dropwise addition of 5 ml Carnoy’s solution (3:1 methanol:acetic acid) with gentle vortexing to avoid clumping. Cells were incubated on ice for 10 min, spun down and supernatant removed. 5 ml of fresh Carnoy’s solution was then added and the cells stored at 4°C. Prior to cytospin, cells were counted and resuspended in fresh Carnoy’s solution to a concentration of approximately 500 cells/µl. 100 ul of cell suspension was then spun onto slides at 10,000 rpm for 2 mins using 3-well adaptors (StatSpin Cytofuge 2). Slides were washed with Carnoy’s solution and allowed to dry. Chromosome specific FISH was then performed with a Texas Red-labeled chromosome 6 alpha satellite probe, following the manufacturer’s protocol (Cytocell).

#### Immunofluorescence

Cells were pre-extracted with ice-cold 1x CSK buffer (10 mM HEPES pH 7.4, 0.3 M sucrose, 100 mM NaCl, 3 mM MgCl_2_) containing 0.1% Triton X-100 for 2 min at room temperature and then fixed with ice-cold 100% methanol at -20°C for 10 min. IF was then performed as previously described (16).

#### Co-Immunoprecipitation (co-IP)

Co-IP was performed as previously described (16). Plasmids used include pcDNA3.1-Flag-CTC1, pcDNA3.1-Flag-STN1 and pCMV6-TEN1 (16,18).

#### Proximity Ligation Assay (PLA)

HeLa 1.2.11 cells were fixed for 20 minutes at room temperature with 4% formaldehyde in 1x PBS followed by permeabilization with 100% methanol for 20 min at -20° C. Subsequent steps were performed with the Duolink PLA kit (Millipore-Sigma) as previously described (24), except the first wash after primary incubation was performed using wash buffer A, not 5% BSA in 1x PBS. Primary antibodies: 1:100 mouse α-STN1, 1:600 rabbit α-SMC3.

#### Image analysis

For IF, FISH and PLA, images were taken on an EVOS FL microscope, using a 40x or 60x objective (Thermo Fisher Scientific). At minimum five images were scored per independent, biological experiment for each condition. Image analysis for Figures 1 and 3D was performed using ImageJ and Figures 2D and 3B with CellProfiler.

#### Supporting information

This article contains supporting information.

## Acknowledgements

We would like to thank Yilin Wang, Margaret Pokalsky and Ross Jaeger for assistance with experiments and Dr. Lauren Ball for assistance with the mass spectrometry analysis. We would also like to thank Dr. Christopher Kasbek, Dr. Carolyn Price and the members of the Stewart lab for helpful discussions. Finally, we thank Dr. Carolyn Price (HCT116 CTC1^F/F^ cells) and Dr. Prasad Jallepalli (Ac-SMC3 antibody) for generously providing cell lines and reagents. Proteomic analysis was performed at the Mass Spectrometry Facility, a University Shared Research Resource at the Medical University of South Carolina, using instrumentation acquired through the NIH shared instrumentation grant program (S10 OD010731).

